# Resolving Protein Conformational Plasticity and Substrate Binding Through the Lens of Machine-Learning

**DOI:** 10.1101/2022.01.07.475334

**Authors:** Navjeet Ahalawat, Jagannath Mondal

**Affiliations:** CCS Haryana Agricultural University, Hisar, Haryana,India; Tata Institute of Fundamental Research, Center for Interdisciplinary sciences, Hyderabad 500046, India

## Abstract

A long-standing target in elucidating the biomolecular recognition process is the identification of binding-competent conformations of the receptor protein. However, protein conformational plasticity and the stochastic nature of the recognition processes often preclude the assignment of a specific protein conformation to an individual ligand-bound pose. In particular, we consider multi-microsecond long Molecular dynamics simulation trajectories of ligand recognition process in solvent-inaccessible cavity of two archtypal systems: L99A mutant of T4 Lysozyme and Cytochrome P450. We first show that if the substrate-recognition occurs via long-lived intermediate, the protein conformations can be automatically classified into substrate-bound and unbound state through an unsupervised dimensionality reduction technique. On the contrary, if the recognition process is mediated by selection of transient protein conformation by the ligand, a clear correspondence between protein conformation and binding-competent macrostates can only be established via a combination of supervised machine learning (ML) and unsupervised dimension reduction approach. In such scenario, we demonstrate that an *a priori* random forest based supervised classification of the simulated trajectories recognition process would help characterize key amino-acid residue-pairs of the protein that are deemed sensitive for ligand binding. A subsequent unsupervised dimensional reduction via time-lagged independent component analysis of the selected residue-pairs would delineate a conformational landscape of protein which is able to demarcate ligand-bound pose from the unbound ones. As a key breakthrough, the ML-based protocol would identify distal protein locations which would be allosterically important for ligand binding and characterise their roles in recognition pathways.

## Introduction

The ubiquitous process of biomolecular recognitions is central to all biological phenomena and has remained an integral part of the research and structure-based drug discovery.^1,2^ With significant upheaval in spatial and temporal resolution in the measurement tools and computer simulation approaches over the last decade, a refined view of underlying atomistic mechanism of the protein-ligand binding phenomena is slowly emerging. ^3^ Due to the periodic upgrades in computer hardwares^4,5^ and GPUs^6^ ligand-recognition in complex solvent-inaccessible cavities of multiple proteins are now regularly being simulated with success.^7–10^ These ligand-recognition simulation trajectories, in combination with the framework of Markov state model (MSM),^11,12^ have served as key resources for an atomic-level characterization of ligand-recognition pathways^9,10,13,14^ and discovery of crucial non-native metastable^13,15^ ligand-bound protein conformations.

However, the access to big data associated with these ‘binding’ simulation trajectories has also raised intrigue if one could distinguish any particular binding-competent protein conformation, that is primed for a specific ligand-bound (native or non-native) pose, over the ligand-unbound protein conformation. ^13^ This question is actually pertinent as establishment of an one-to-one correspondence between a specific protein conformation, that is pre-selected for a complementary ligand-protein complex macro state, would assign a structural/functional connotation between these two. However, until recently no such precedence exists. The inherent issue of being unable to reconcile conformational heterogeneity prevalent in protein/ligand recognition project is often known to stem from the dynamical protein fluctuations associated with overwhelming big MD-derived trajectories. Biomolecular simulations are intrinsically multidimensional and generate noisy trajectory sets of ever-increasing size. A potential solution in this aspect would be a suitable state-space decomposition of protein conformations via dimensional reduction and identifying its ligand-binding-competence. However, recent investigations have shown that an automated assignment of a given protein conformation to a complementary native or non-native ligand-bound pose is extremely challenging, if not impossible. This has given rise to the theme of conformational plasticity in protein.^13,16–18^ Figure 1 provides a schematic illustration of the theme of conformational plasicity.

**Figure 1:**
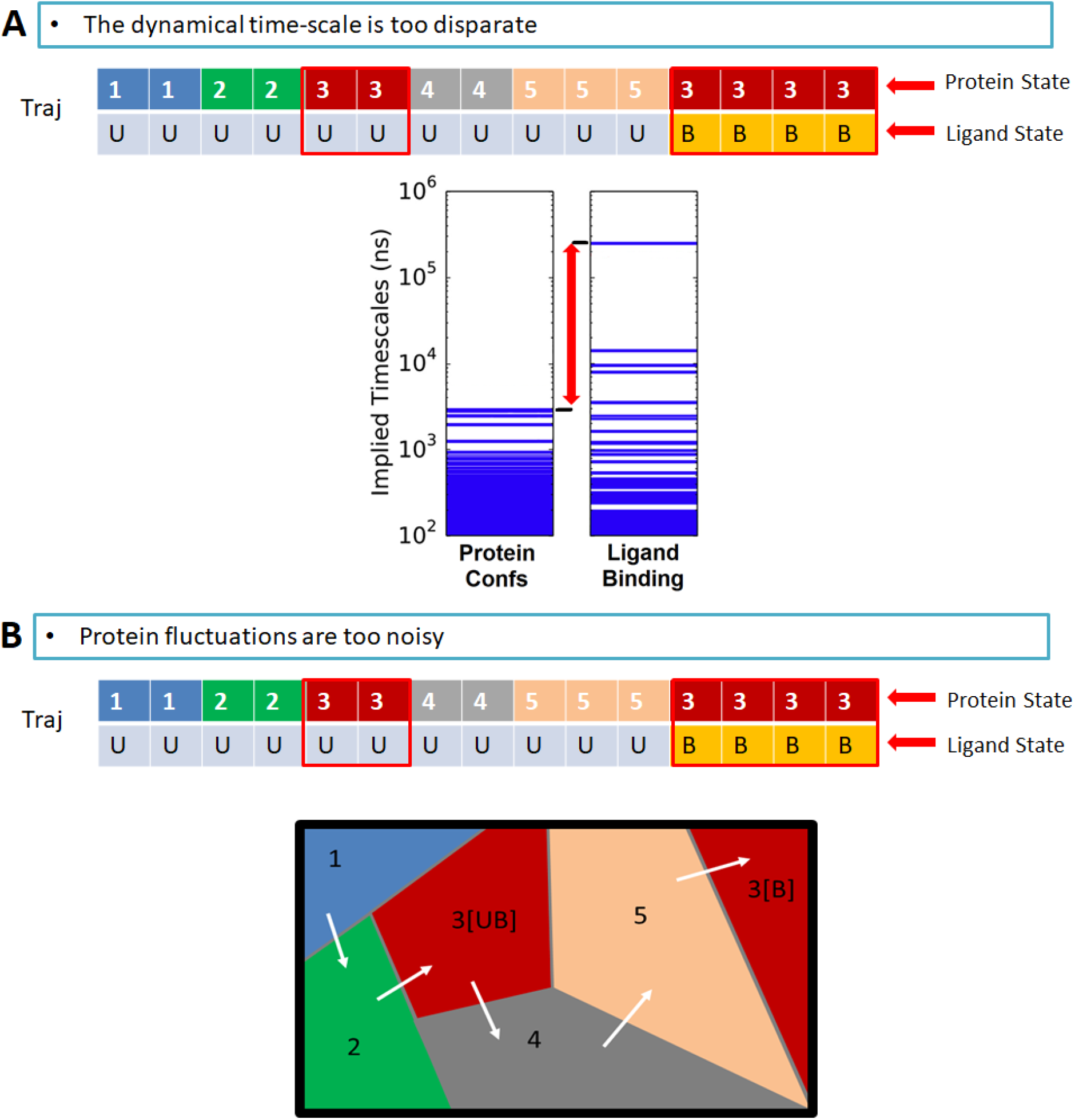
Zooming into Possible origins of protein-conformational plasicity. Two possible reasons are illustrated a) The dynamical time-scales of protein conformational fluctuations and ligand binding event are too disparate, b) The Protein conformational fluctuations are too noisy to truely resolve ligand bound states from the unbound ones.

For example, a recent attempt at correlating binding-competent states to distinct protein conformation had resulted in ‘mixing’ or ‘co-existence’ of multiple protein conformations. ^13^ The investigation eventually required subsequent manual, iterative intervention in combination with kinetic clustering of the MD trajectories to reconcile conformational plasticity and ligand binding. We surmise that there might be possibly two reasons for the inability to assign ligand-bound and unbound states to individual protein conformation (figure 1):

- either the dynamical time-scale of protein conformational change and ligand-binding event is too disparate to expect that for a specific protein conformation, a ligand-bound pose can exist.
- or, the underlying stochasticities of MD simulation trajectories and rapid fluctuations in the protein conformations during the period of the simulation deter an establishment of correspondence between protein conformation and ligand location around the protein.

In the present work, we propose to alleviate the second issue via machine learning (ML) approach, a technique that has garnered its well-deserved attention in biomolecular sciences due its ability to process large-scale data^19^ in an efficient manner. In particular, we demonstrate that a judicious integration of a popular supervised ML classifier, namely ‘Random Forest (RF) algorithm’, with an unsupervised dimension reduction technique, namely, time-structured independent component analysis (TICA)^20–22^ would enable non-overlapping assignment of binding-competent protein conformations. As a case study, we analyze a set of multi-microsecond long MD simulation trajectories of ligand-binding event in two proteins, namely L99A mutant of T4 Lysozyme and cytochrome P450. We first show that for cytochrome P450, which recognises the substrate via an induced-fit based mechanism,^9^ the protein conformations can be automatically classified into substrate-bound and unbound state though an unsupervised dimensionality reduction technique. However, for system such as L99A T4 Lysozyme, in which the recognition process is mediated by selection of transient protein conformation, a clear correspondence between protein conformation and binding-competent macrostates can only be established via an intervention of supervised machine learning and unsupervised dimension reduction approach.

## Results and Discussion

Archtypal proteins Cytochrome P450 and L99A T4 Lysozyme, in their act of substrate recognition, serve as the systems of our current investigation. Both the proteins have remained the subject of multitudes of precedent experiments and simulations,^9,23–28^ due to their intriguing features of ligand recognition ability in an otherwise deeply buried cavity. In particular, recently reported multi-microsecond long unbiased Molecular Dynamics (MD) simulation trajectories in both these systems by our group^9,10^ have captured the complete binding processes of substrate from solvent to the buried cavity of these two proteins, as demonstrated via the respective ligand-binding time profiles in both these systems in figure 2. Figure 2A represents three independent MD trajectories of successful substrate recognition in cytochrome P450 in terms of the time profiles of distance between the centre of mass of the ligand and that of the designated cavity located near the heme active site of the protein. On the other hand, figure 2B depicts the same for L99A T4 Lysozyme In both the systems, in each of the three representative trajectories, the eventual simulated bound pose came within an angstrom-level accuracy with that of the crystallographic bound pose, attesting to a precise capture of the ligand recognition event in real time simulation. However, detailed analysis of binding trajectories in these two systems had previously revealed distinct and contrasting recognition mechanisms: The camphor recognition in cytochrome P450 was found to take place via a single dominant pathway which is mediated by an induced fit mechanism^9,28^ resulting into a long-lived on-pathway intermediate. On the contrary, ligand binding trajectories in L99A T4 Lysozyme^10^ further revealed multiple distinct pathways of ligand recognition via subtle fluctuation of the helices around the binding cavity, prompting the ligand to select certain kinetically transient protein conformations in a bid to find the cavity.

**Figure 2:**
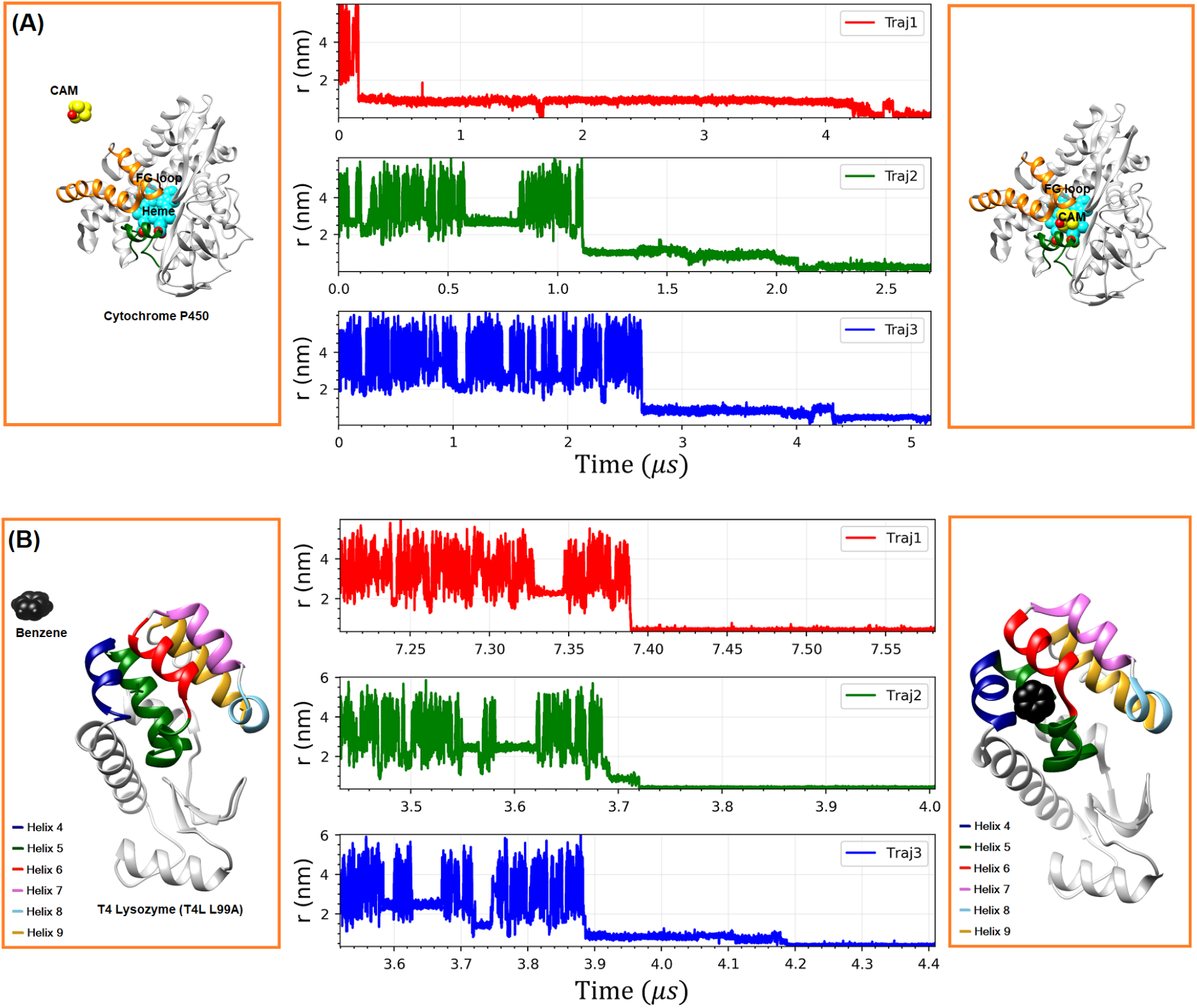
Time profile of cavity-ligand distances as analyzed from three independent simulation trajectories of A. Cytochrome P450 and B. L99A T4 Lysozyme. Also shown are the simulated unbound and ligand-bound pose of protein.

The availability of multiple set of ligand-binding simulation trajectories in both these systems (with contrasting recognition mechanism), recorded at a frequent time-interval, prompted us to identify the key protein conformations in each of these trajectories and more importantly, to investigate if these protein conformations can individually be assigned to any particular ligand-binding-competent macrostate.

Towards this end, first we focussed our interest on substrate-recognition process in cytochrome P450, with the aim of resolving binding-competent protein conformation from that of unbound state. To characterize the key protein conformations in these dynamically evolving simulation trajectories, we first determined the minimum distances among all the heavy atoms between each residue-pairs and curated these pair-wise distances for all time-series in the form of a large matrix. Subsequently, we performed a dimensional reduction of these contact features using time-structured independent component analysis (TICA)^20,21^ (see figure 3, left, for protocol). Widely perceived as a popular unsupervised dimension reduction technique, TICA is known to project the time-series data along the direction, along which the time correlation is maximized and hence would produce dynamically slowest projection. Figure S1 A) projects three independent simulation trajectories of cytochrome P450 in a free-energy surface (FES) along the two most slowest time-structured independent component (TIC) dimensions. We find that the projection of the protein conformations in the FES represents one or more local minima, suggesting that TICA is able to identify key protein conformational sub-states.

**Figure 3:**
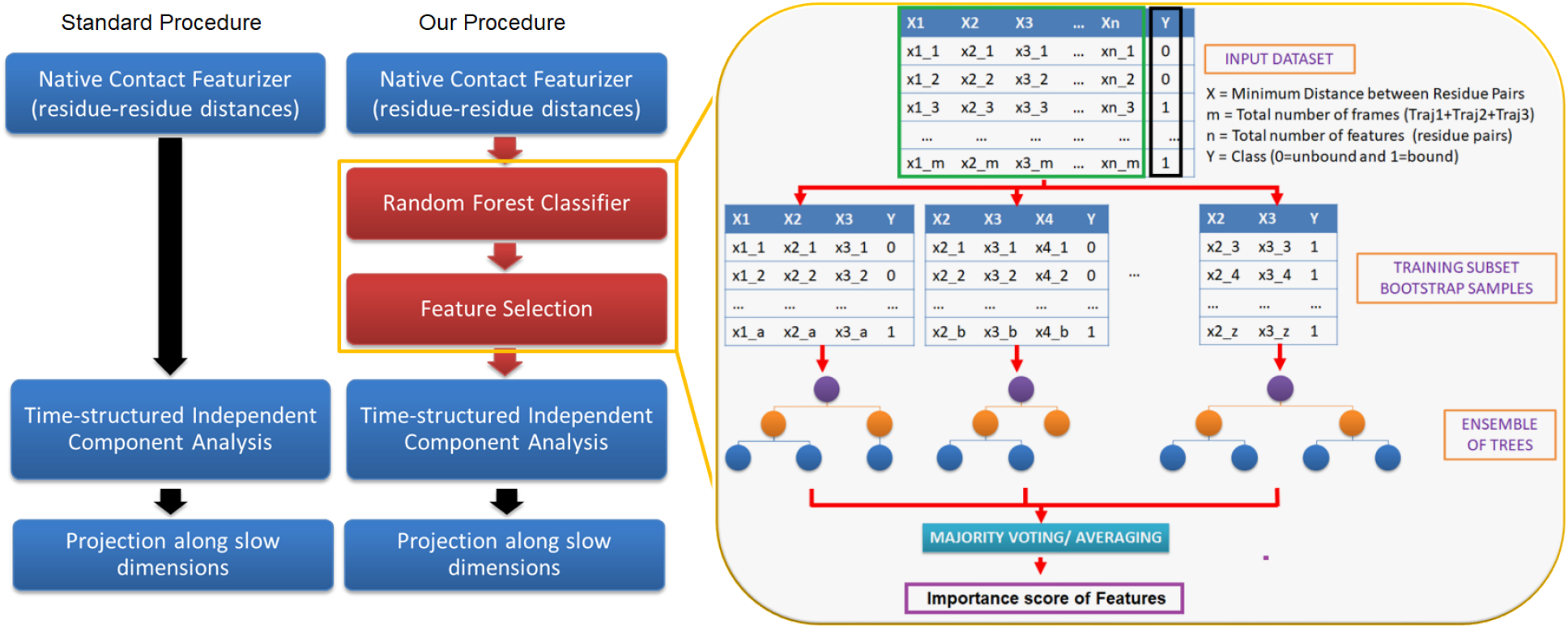
Illustration of proposed scheme involving random forest protocol. Left scheme highlights the standard procedure of dimensionality reduction of protein conformation via TICA. ^20,21^ Right scheme demonstrates our proposed scheme of state-space decomposition via combining random-forest based supervised learning with TICA.

Next, in order to probe if these protein conformations of cytochrome P450 can be ascribed to specific binding-competent states, we annotated the respective ligand locations on the dimensionally reduced protein conformational landscape. To this end, we calculated the average cavity-ligand distance for the conformations of each grid corresponding to the FES obtained from TICA analysis. Figure 4 depicts the overlay of average cavity-ligand distance on TICA-derived projections of protein conformation. As evident, the assignment of ligand-bound poses (blue dots) and ligand-unbound poses (red dots) are clearly non-overlapping across all three trajectories, suggesting that the unsupervised assignment of ligand-location on conformational landscape, projected on the slowest TICA dimensions, is able to distinguish the ligand-bound state from the ligand-unbound state. A possible reason for clear segregation of substrate-bound protein conformation from that of unbound conformation in a dimenisionally reduced subspace might be that the recognition process in cytochrome P450 takes place via a single dominant pathway in which the substrate induces a long-lived intermediate before getting bound in the heme pocket. The long time spent by the ligand in a specific protein conformation allows a clear separation of time-scale of substrate-bound protein conformations from that of unbound conformation.

**Figure 4:**
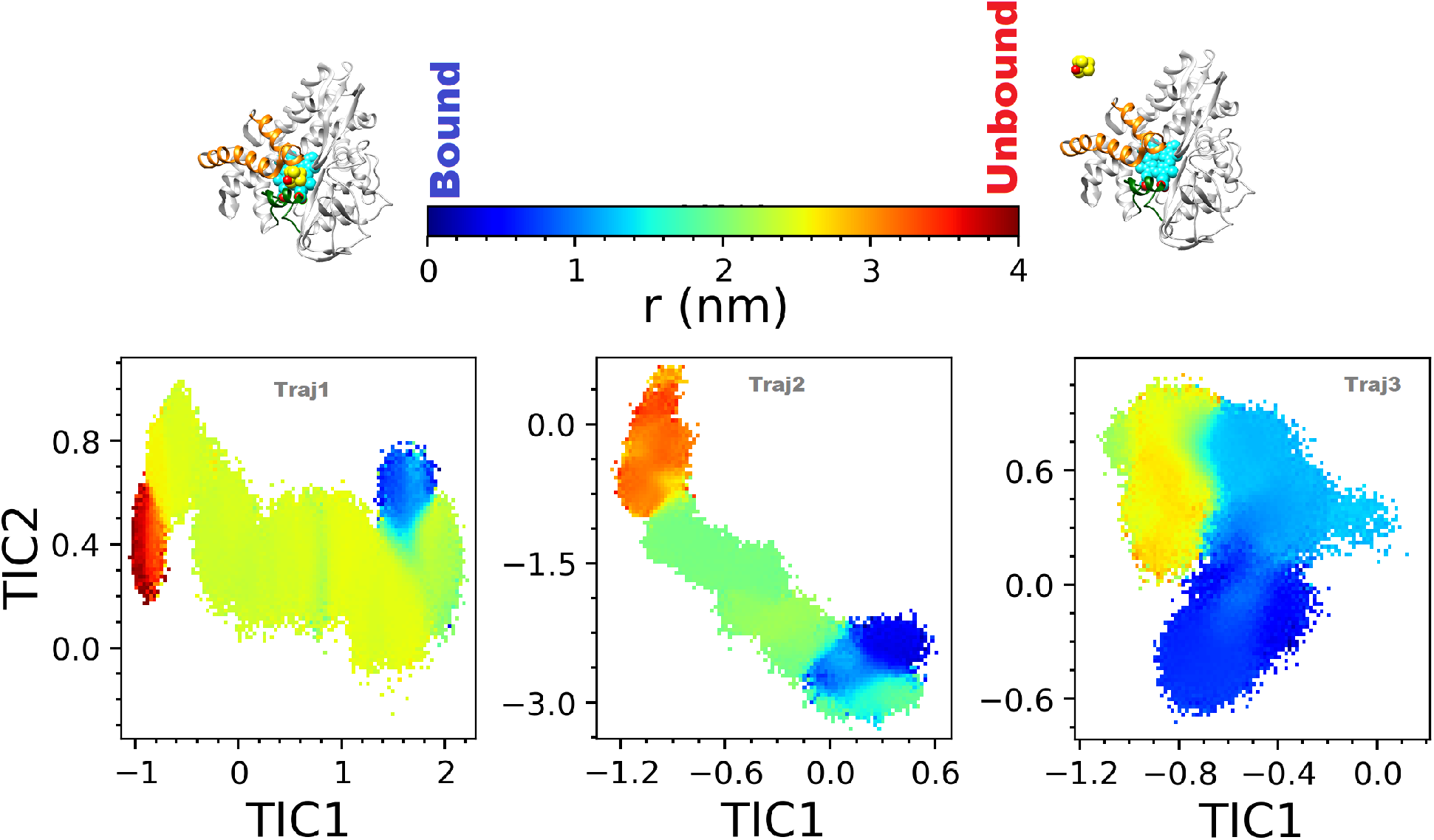
The scatter plot of the cytochrome P450 cavity-ligand distance overlayed on the FES along two slowest Time-lagged independent component of the protein motion.

Encouraged by the ability of TICA-based dimension-reduction approach in distinguishing binding-competent protein conformations in low-dimensional conformational landscape of cytochrome P450, we wanted to explore if the same can be achieved in another system namely, benzene binding to L99A T4 Lysozyme. As described before, previous atomistic simulations^10^ had demonstrated that the ligand binding mechanism in L99A T4 Lysozyme involves multiple ligand-binding pathways through selection of transient conformations, as opposed to an induced fit binding in cytochrome P450. Hence we thought if sim ilar unsupervised dimensionality reduction approach would work in this system.

Similar to the protocol described in case of cytochrome P450 (figure 3,left, standard procesure), we first projected three independent simulation trajectories of L99A T4 Lysozyme in a free-energy surface (FES) along the two slowest time-structured independent component (TIC) dimensions (figure S1 B). We find that, similar to cytochrome P450, TICA is able to identify multiple basins of protein conformations. However, our attempt to annotate the protein conformational landscape by ligand location showed that in majority of the trajecories (in two out of three), the ligand-bound and unbound conformations are considerably mixed, suggesting that this unsupervised dimension reduction approach is not able to distinguish ligand-bound state from that of unbound state (figure 5). This demonstrates a classic case of conformational plasticity that is inherent in T4 Lysozyme, as previously examplified by subtle helix movement (as introduced in figure 1)

**Figure 5:**
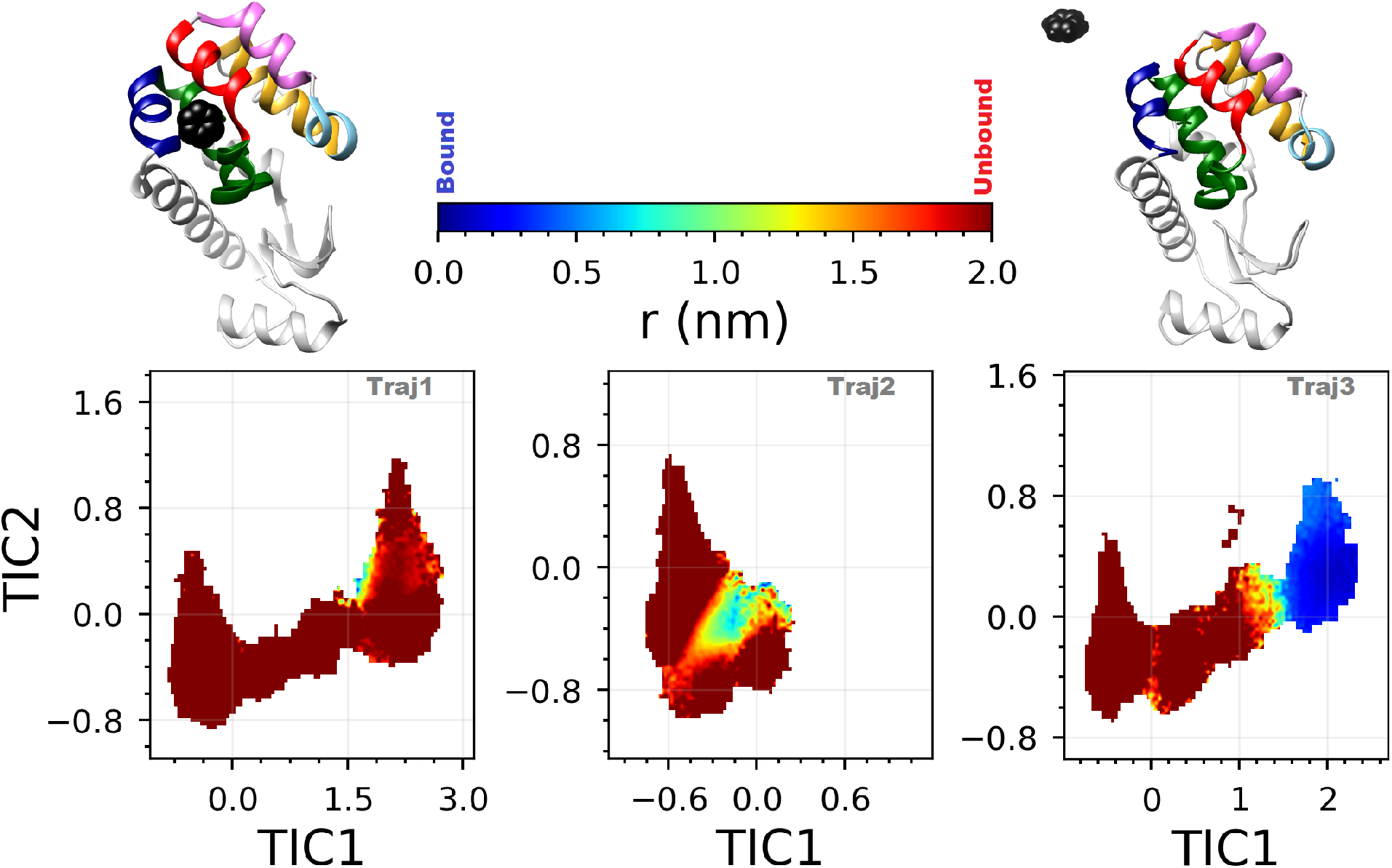
Free energetics of Protein conformational subspace of three T4 Lysozyme/benzene simulation trajectories along top two TIC dimensions which were derived using all residue pairs’ distances of the protein. The free energy surfaces (FESs) are colored according to the T4 Lysozyme cavity/ligand (average) distance.

An analysis of implied time-scale of protein conformational fluctuation and ligand binding dynamics in T4 Lysozyme revealed that these two time-scales are not significantly different from each other (see figure 1 for illustration), with the slowest time scales ranging around 10^3^ nanosecond for both processes.(Figure 6) The overall analysis indicates that the inability in finding a correspondence between protein conformation and its ligand-binding competence in this particular system of L99A T4 Lysozyme/benzene is not due to the large difference in time-scale: rather the automated, unsupervised dimension reduction technique is not able to assign or identify corresponding protein conformation for a ligand-bound (and ligand-unbound) states. Accordingly, we postulated if a supervised machine-learning approach can be *a priori* introduced to refine the contact-featurizers of the simulation trajectories, which can then be subsequently subjected to an unsupervised machine-learning technique.

**Figure 6:**
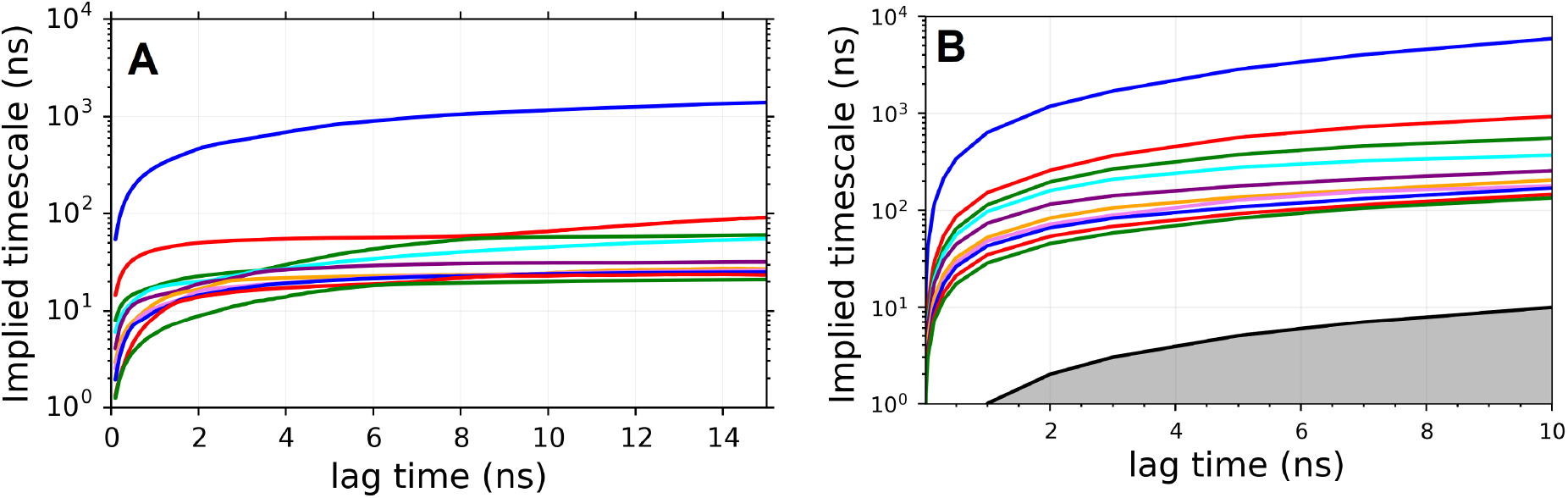
Implied time-scale of A) Protein/ligand binding kinetic and B) Protein conformational kinetics

Towards this end, we employed a popular supervised machine learning approach called ‘Random-forest (RF) algorithm’,^29–31^ (see figure 3, right for proposed protocol) to rank-order the residue-pairs which has correlation with the ligand binding. Recent reports^32–34^ of promising performance by RF as a supervised classifier of biomolecular simulation data as well as chemical sciences^35^ prompted its choice in the present work. RF is a powerful machine-learning as well as feature selection method which can directly identify a subset of useful feature from a large set of input variables. In addition to feature selection, it also has provisions for ranking the selected features according to their relevance for predicting the output. Several such importance measures have been proposed in the literature. As a part of the implementation of RF algorithm in current investigation, all trajectory frames are divided into two states (‘bound’ and ‘unbound’) using distance cut-off of 0.5 nm between ligand and binding cavity. Next, native contacts (based on the crystal structure of L99A T4 Lysozyme) were used to identify the residue pairs and minimum distance between the heavy atoms of residue pairs were invoked as input dataset in RF algorithm. Here, we employed default method of RF algorithm, as implemented in scikit-learn python package^36^ which computes variable importance as the mean decrease in impurity (‘Gini importance’)^31^ (see Method for details). The algorithm identified top 200 residue-pairs based on their ‘importance to the ligand binding’. We then proceeded with RF-ranked distance pairs as inputs for subsequent TICA-based projection of protein conformation. Very interestingly, the annotation of the cavity-ligand distances on these newly obtained FES (figure 7) showed that ligand-bound (blue dots) and ligand-unbound (red dots) states are mutually segregated in the reduced dimension underlying all three trajectories. This implies that the *a priori* supervised filtering by a RF classifier has effectively assisted in refining the projection of the protein conformation such that assignment of ligand-bound and ligand-unbound macro states is feasible in all three trajectories.

**Figure 7:**
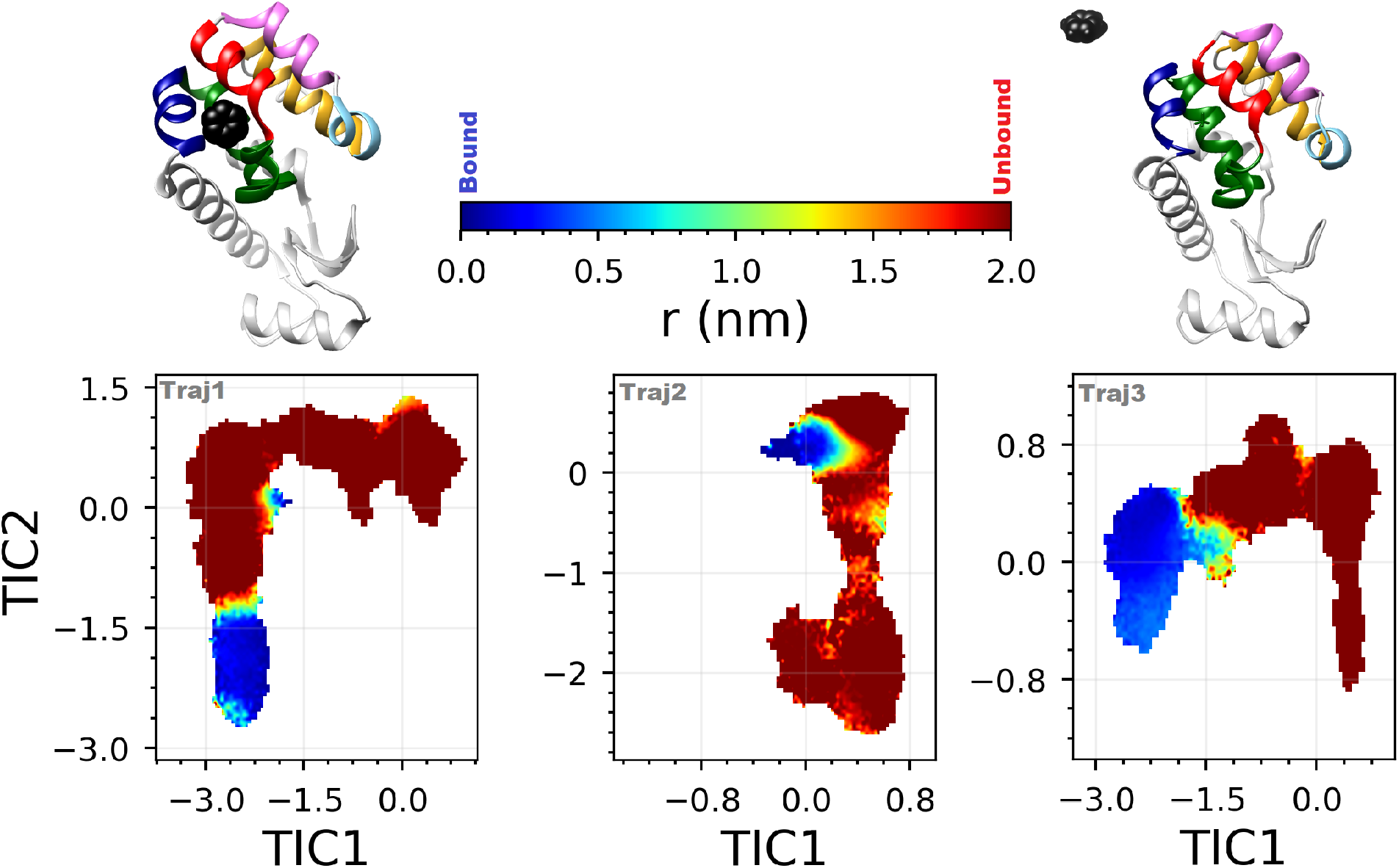
Free energetics of Protein conformational subspace of three T4 Lysozyme/benzene simulation trajectories along top two TIC dimensions which were derived using Random-forest ranked 200 residue pairs’ distances of the protein. The free energy surfaces (FESs) are colored according to the T4 Lysozyme cavity/ligand (average) distance.

A key output of RF classifier is an ‘importance or sensitivity’ score that it assigns to each of the residue pairs (features) based on its correlation to the ligand binding. RF classifier has previously been used for identifying protein allostery.^34^ Annotation of the ‘important’ residues, as predicted by RF classifier (figure 8, bottom), on the three-dimensional cartoon representation of the L99A T4 Lysozyme, indicates that many of these residue-pairs are located in proximity to the designated binding site of this protein. In particular, ligand entry in these trajectories are known to occur on the helices around the C-terminal domain of the protein and designation of ‘high importance’ by RF classifier to certain residue pairs around C-terminal domain is expected. However, the classifier also predicts a set of residues which are distal from binding pocket (N-terminal domain of the protein) and yet are considered ‘important’ by RF classifier towards the cause of ligand binding. Intriguingly, some of these distal pairs (involving residues 18, 22, 137) have previously been reported to be allosterically important by other experimental^37^ as well as theoretical^38^ studies.

**Figure 8:**
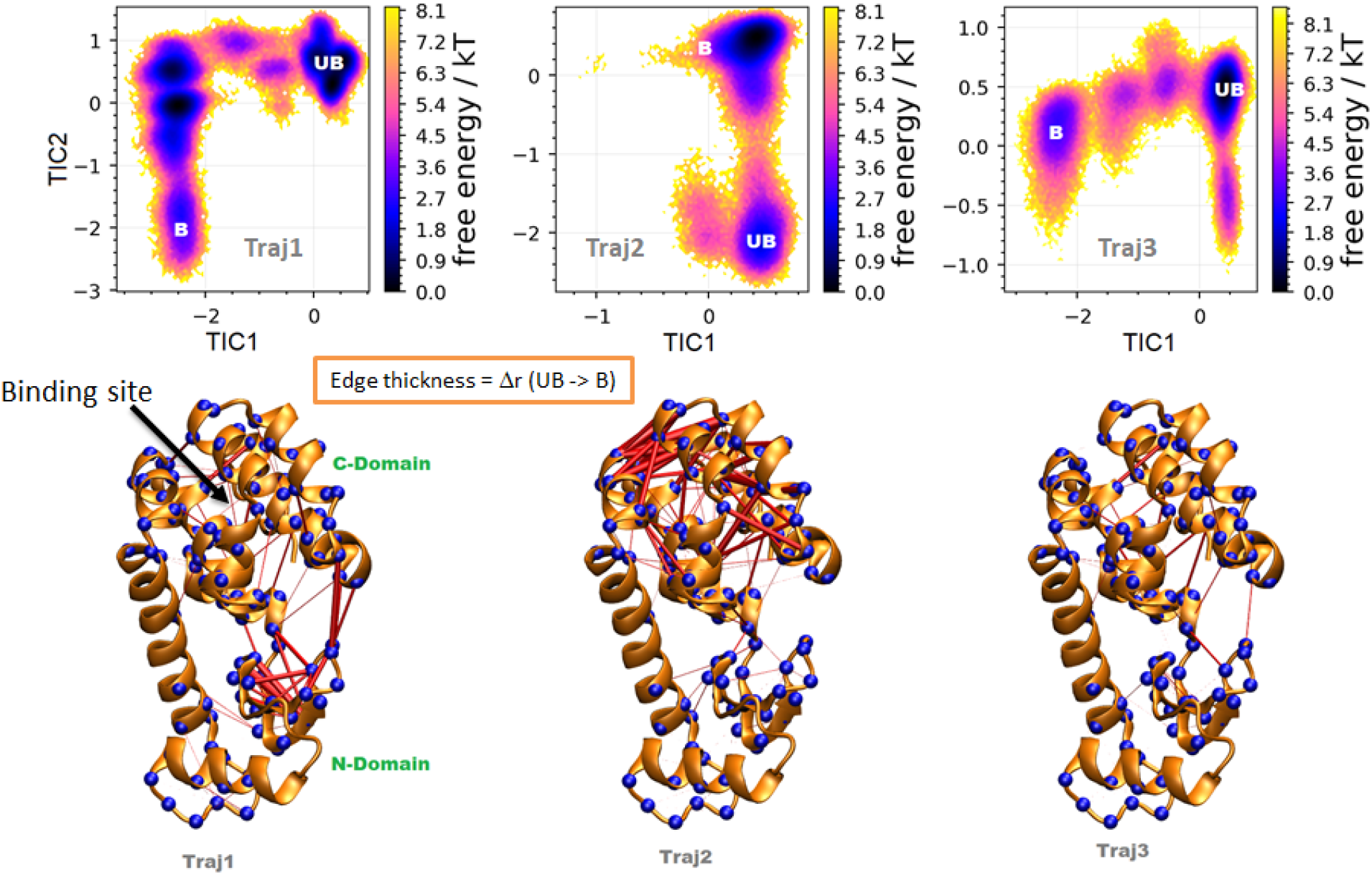
Difference in inter-residue network between ligand-bound and Unbound conformations of T4 Lysozyme. Trajectory 1 and trajectory 2 show significant changes in distal residue networks upon ligand binding while trajectory 3 involves uniform changes in all protein locations.

A closer look of the TICA projection of conformational landscape for each of three trajectories, obtained after the supervised learning of random forest featuriser, (figure 8) indicated that the underlying FES are distinct in each cases, while clearly demarcating the ligand-bound and un-bound (labelled ‘B’ and ‘UB’ respectively in FES) states in spatially resolved locations of the FES. Very interestingly, past investigations had reported three different ligand-entry pathways for these three trajectories. To understand the role of all selected residues in the binding process we calculated the change in average fluctuation of residue distances on shifting from unbound state to bound state. Here, we have generated the ensemble of randomly selected protein conformations from bound and unbound regions (marked ‘B’ and ‘UB’) of the FES shown in figure 8,top, for each trajectory. To visualize the difference in the bound and the unbound states, the changes in average fluctuation between these selected residue pairs were projected on the 3D structure in the form of networks (see red lines). The thickness of a connection represents the average difference in inter-residue distances between ligand-bound (B) and unbound (UB) conformations of T4 Lysozyme. Interestingly, each trajectory shows a different pattern of connection between the selected residue pairs, which might be partly due to their mutually distinct ligand-recognition pathways. In trajectory 1, the binding of ligand is known to occur through the helix 4 and helix 6 but the thickness of residue pairs is higher for interfacial residue-pairs of N-domain and C-domain as well as intra N-terminal domain indicating possible allosteric role of these distal residues in the binding process. Trajectory 2 shows the large conformational fluctuations near binding site (C-Domain), while Trajectory 3 involves uniform changes in all protein locations.

## Conclusion

In summary, the current investigation put forward a machine-learning based recipe to efficiently distinguish binding-competent (native or non-native) protein conformation from that of ligand-unbound state. By considering the example of ligand recognition by cytochrome P450 and L99A T4 Lysozyme as two interesting case-studies, we demonstrate an efficient combination of supervised and unsupervised machine-learning approach can resolve inherent protein motion from that of ligand motion. We demonstrated that when the substrate-recognition is guided by an induced-fit mechanism which is accompanied by a long-lived intermediate (as in case of cytochrome P450/camphor binding), an automated separation of ligand-bound and unbound protein conformation is plausible in a dimenisonally reduced sub-space. However, for cases where recognition process is guided by selection of transient protein conformation by ligand (as in case of T4 Lysozyme/benzene binding), an unsupervised approach might fail. In such scenario, when combined with supervised classification algorithm, namely RF classifier, an unsupervised dimension-reduction technique, such as TICA, is able to individually assign ligand-bound and unbound state to distinct protein conformational state-space. Apart from identifying the important amino-acid residue-pairs proximal to the binding cavity of the protein, the protocol also attests importance to multiple residue-pairs which are distal from the ligand-recognizing cavity, thereby discovering allosteric importance of many amino-acid residues in ligand-binding.

Sheer volume of the data currently being generated by molecular simulations poses serious challenges for analysis and interpretation. Accordingly, recent times have seen employment of many tools inspired from ML methods, most notatably, neural network based nonlinear models,^39–43^ the unsupervised nature of majority of this ML tools are often criticised for obstruction to human-interpretable insights. As a result,judicious introduction of supervised classifiers within ML based protocols are gaining tractions in biomolecular and chemical sciences.^32,33,44–46^ Towards this end, the current investigation offers an effective solution in resolving underlying mechanism of ligand binding in conformationally plastic receptors^16–18^

## Method and Model

The previously reported multi-microsecond long MD simulation trajectories by Mondal et. al.^10^ had described the recognition processes of benzene to L99A mutant of T4 Lysozyme protein at an atomistic precision. Three MD trajectories (simulation length ranging between 4-7.5 micro second) from the previous investigation form the base of the current investigation. The simulation model and methods have been detailed in the work by Mondal et al.^10^ Briefly, the protein and benzene were modelled by charmm36 all-atom force fields^47^ and simulated in explicit presence of TIP3P water^48^ and ions in periodic box. The MD simulation trajectories is unbiased and unguided in nature, with no restriction on the protein and ligand movements. A NPT ensemble has been adopted with average temperature of 300 K and pressure of 1 bar. The simulations employ a time-step of 2 femtosecond and the coordinates had been saved at an interval of 10 picosecond.

The RF classification algorithm, ^29–31^ an ensemble learning and supervised machine learning method was first proposed by Breiman.^29^ RF is one of the most powerful learning as well as feature selection method which can directly identify a subset of useful feature to solve a problem from a large set of input variables. In RF algorithm, *n* samples from the input training dataset were drawn as a training subset to generate an ensemble of tress (a decision tree for each subset). These randomly selected multiple unrelated decision trees establishes the random forest or random decision forest. From this ensemble of tress an optimal classification result is obtained by averaging method or by the voting method. A representative flowchart of the RF algorithm is shown in Fig. 2,right.

In addition to feature selection, it also allows to further rank the selected according to their relevance for predicting the output. Several such importance measures have been proposed in the literature. All these measures of importance provide interesting alternatives to attribute ranking based on the (adjusted) p-values obtained from classical statistical tests. The main advantage of these measures with respect to these statistical tests is that they do not make any assumption about the problem (such as gaussianity, linearity, or independence) and they are potentially able to detect multivariate effects, i.e. attributes that are only relevant through interaction with others. However, the tree-based importance measures are not yet as well principled as statistical tests, because their limitations and biases are not yet fully characterised, although research in this area is ongoing.

The random forest analysis is set up according to the following procedure:

- All the MD frames are divided into two states (bound and unbound) using distance cut-off 0.6 nm between benzene and cavity for T4 Lysozyme.
- Native contacts (based on the crystal structure) were used to identify the residue pairs and minimum distance between the heavy atoms of residue pairs were used as input dataset in RF algorithm.
- Here, we employed default method implemented in scikit-learn python package^36^ which compute variable importance as the mean decrease in impurity of gini impurity.^31^ Random forest algorithm implemented in sklearn 0.22 was employed with *n*_*estimators*_=1000, bootstrap=True, and the features were considered by using the Gini feature selection.

In this work, we employ an unsupervised dimension-reduction technique, namely TICA, which has garnered popularity for its ability to project the MD simulation data along kinetically slowest projection.^20,21^ The method has its origin in the independent component analysis (ICA), a statistical and computational technique which transforms a multidimensional data into statistically independent components and is very popular in the field of signal processing. As detailed elsewhere, ^20,21^ TICA identifies the slowest coordinates without losing important kinetic information via maximizing the autocorrelation function of the projection of the simulated data for a given lag-time. Here, for both the systems of cytochrome P450 and L99A T4 Lysozyme, we have used time series data of residue-residue distances from the simulated trajectories as input for TICA with 10 ns lag-time. For visualization purpose, the trajectories were projected along the first two slowest dimensions (TIC1 and TIC2). The inter-residue distance was defined as the minimal distance between heavy atoms of that residue pair.

## Supplemental Information

Figure with Free energy surface of Protein conformation along slowest TICA-derived dimensions.

## Acknowledgements

This work was supported by computing resources obtained from shared facility of TIFR Centre for Interdisciplinary Sciences, India. JM would like to acknowledge research intramural research grants obtained from TIFR, DAE, India, Ramanujan Fellowship and Core Research grants provided by the Department of Science and Technology (DST) of India (CRG/2019/001219).

