## Supplementary figures for "Resolving Protein Conformational Plasticity and Substrate Binding Through the Lens of Machine-Learning"

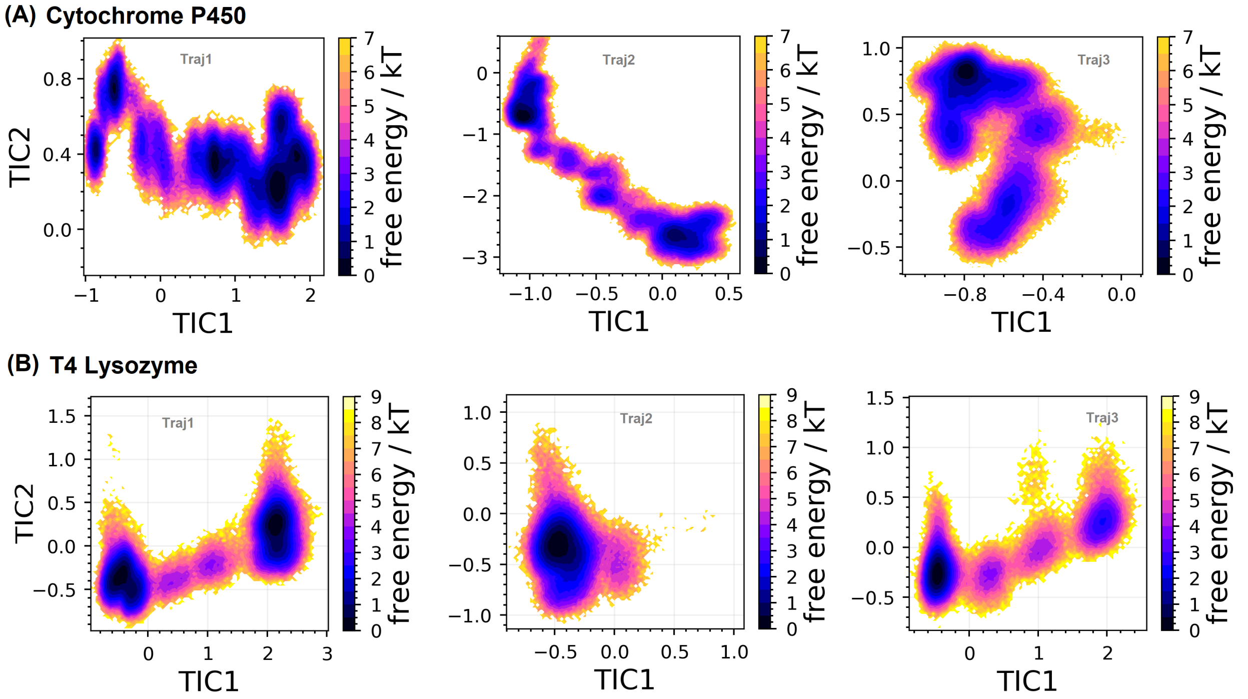


**Figure S1:** Conformational Free energy landscape for A)cytochrome P450 and B)T4 Lysozyme along slowest TICA-derived dimensions.
